# Ferumoxytol dynamic contrast-enhanced MRI for *in vivo* longitudinal cotyledon perfusion assessment with pathology correlation in a rhesus macaque thrombotic injury model

**DOI:** 10.64898/2026.08.04.742075

**Authors:** Ruo-Yu Liu, Logan T. Keding, Ruiming Edmondson, Jessica Vazquez, Kathleen M. Antony, Kevin M. Johnson, Dinesh M. Shah, Thaddeus G. Golos, Aleksandar K. Stanic, Oliver Wieben

**Author notes:** **<u>Corresponding author:</u>** Oliver Wieben 1111 Highland Avenue, Suite 1127 Wisconsin Institutes for Medical Research UW-Medical Physics and Radiology Madison, WI 53705-2275.

## Abstract

**Introduction:** While placental perfusion and pathology jointly affect pregnancy outcomes, cotyledon-specific perfusion across gestation and its correlation with local injury is not yet well understood. Ferumoxytol dynamic contrast-enhanced magnetic resonance imaging (DCE-MRI) offers a promising way to noninvasively identify cotyledons across gestation and quantify longitudinal cotyledon-specific perfusion changes. Additionally, intraplacental injection of bioactive fibrin sealant allows us to model thrombotic placental injury and further assess cotyledon-level relationships between perfusion and significant injury.

**Methods:** Pregnant rhesus macaques (N=13) received intrauterine saline or fibrin sealant injections at gestational day (GD) ∼101 and underwent ferumoxytol DCE-MRI at GDs ∼100, 115, and 145. Placental perfusion domains derived from contrast arrival time were segmented at each imaging time point and matched to cotyledons identified following tissue collection by cesarean section, with cotyledon perfusion quantified longitudinally and correlated with cotyledon-specific quantitative histopathology.

**Results:** All pregnancies were successfully carried to term. Fibrin sealant injections induced significantly higher levels of placental pathology compared to saline controls. MRI-derived perfusion domains were largely consistent across gestation and showed predominantly one-to-one correspondence with term cotyledons, with successful perfusion-pathology pairing achieved in 153 cotyledons. Longitudinal cotyledon perfusion changes showed significant positive correlations with villous agglutination injuries.

**Conclusions:** Feasibility of noninvasively tracking placental cotyledon perfusion using ferumoxytol DCE-MRI was demonstrated, and the efficacy of the rhesus macaque thrombotic injury model was confirmed. The positive perfusion-pathology correlations suggested intrinsic placental regulatory mechanisms and functional plasticity. This new framework is promising for future translational studies and validation of *ex vivo* cotyledon perfusion models.

**Highlights:**

- Longitudinal tracking of placental perfusion domains with ferumoxytol MRI
- Successful matching of cotyledons and MRI-derived perfusion domains
- Confirmed thrombotic injury-model induced cotyledon pathology
- Maternal perfusion compensation in presence of villous pathology

## Introduction

The placenta is a temporary organ that constitutes the maternal-fetal interface (MFI) of pregnancy, governing the transport of vital resources, such as nutrients, oxygen, hormones, and pharmaceuticals. Connected to the fetus via the umbilical cord and the mother via decidual tissue, the placenta contains anatomically separate maternal and fetal circulatory systems and facilitates material exchanges between the two.

Placental vasculature is specialized to ensure sufficient maternal-fetal transfer and actively changes morphologically and physiologically during gestation. Oxygenated maternal blood fills the intervillous space (IVS) through remodeled decidual spiral arteries, bathing chorionic villi and allowing molecular exchanges without mixing maternal and fetal blood. Placental septa originating from the basal plate project toward the chorionic plate, completely or partially dividing the IVS into individual perfused regions, termed cotyledons. These compartments can contain one or more spiral artery entries and maternal veins [1], enabling the full cycle of maternal blood entry and return within a single cotyledon. Therefore, cotyledons can be conceptualized as the functional units of perfusion in the placenta, with maternal-fetal transfer being directly affected by maternal and fetal blood inflow/outflow and syncytial permeability. The development and health of the fetus are thus largely influenced by placental regulation of uteroplacental perfusion at the MFI.

*Ex vivo* dual perfusion models of single human placental cotyledons have been one of the main approaches to study the structural and functional significance of cotyledons in placental perfusion. Panigel et al. introduced the concept of mimicking *in vivo* placental perfusion in a laboratory setting [2], with Schneider et al. later simplifying the approach by inserting cannulae into the IVS instead of a spiral artery [3]. By preserving the fine vascular structure of isolated cotyledons of delivered placentae, this model imitates *in vivo* flow conditions of both maternal and fetal compartments and has been used for estimating exchange rates of a wide range of molecules at the MFI [4]. In addition, recent advances in *ex vivo* experimental modeling are aiming to both closer resemble *in vivo* physiological conditions and standardize the methodology [5].

One of the most critical inputs to this model is the flow rate of maternal blood influx into the placental IVS. *In vivo* measurements of blood flow with Doppler ultrasound throughout the placenta have been proven difficult due to various spiral artery orientations, location-dependent spiral artery inflow, and largely nonlinear flow trajectories [6–8]. Other methods for *in vivo* flow and perfusion quantification typically require applications of exogenous imaging markers, with the major contenders being X-ray angiography with iodine contrast, positron emission tomography (PET) with oxygen-15-marked water molecules, and dynamic contrast-enhanced magnetic resonance imaging (DCE-MRI). However, X-ray angiography and PET without justifiable medical benefits to pregnant patients are generally avoided due to the potential negative effects of ionizing radiation on maternal health and fetal development [9,10]. DCE-MRI, while not involving ionizing radiation, often utilizes Gadolinium-based contrast agents (GBCAs), which have been shown to pass the placental barrier and enter fetal circulation in mice [11] and nonhuman primates [12] and thus raise concerns about their yet unclear long-term effects [13].

Recently, the off-label use of ferumoxytol as an MRI contrast agent has been proposed. Ferumoxytol is a superparamagnetic iron oxide agent with an FDA-approved indication for intravenous treatment of iron-deficiency [14,15], demonstrating good effects and tolerance in patients including pregnant women [16,17] and pediatric patients [18,19]. Ferumoxytol-enhanced MRI has been shown helpful in assisting diagnoses of placenta accreta spectrum [20–22] and contrast-enhanced functional imaging in the placenta [23–25], with its safety assessed in both nonhuman primates [26,27] and humans [28,29]. Ferumoxytol DCE-MRI is hence a favorable option to measure maternal blood flow in the placenta.

Another challenge for the in vivo validation of the ex vivo perfusion model parameters is measuring maternal perfusion in the placenta both at the cotyledon level and across different gestational stages. While placental perfusion measurements in animals using DCE-MRI have been performed in several studies on the whole-organ level [30–35], discerning individual contributions of each cotyledon remains generally challenging due to the difficulty of identifying cotyledons on MR images. Frias et al. thus [36] proposed a workflow to identify perfusion domains based on spiral artery entries with Gadolinium DCE-MRI and a generalized impulse response model [37], which demonstrated promising spatial resemblance between MRI-derived perfusion domains and cotyledons in rhesus macaque placentae.

Our group has then adapted and modified this workflow to measure cotyledon-specific perfusion in several studies using Ferumoxytol [38–42], showing good spatial agreements between patterns of contrast agent inflow and cotyledons observed in delivered placental tissue. Ferumoxytol, with its prolonged intravascular half-life (14- 15 hours) [43], allows easier contrast arrival time calculation using simple sigmoid fits and eliminates the need to specify arterial input functions. Furthermore, its larger molecular size compared to GBCAs likely warrants lower permeability across placental septa and therefore better contrast between septum-separated cotyledons. This technique thus provides a promising way to label individual cotyledons and track their structural and functional changes over gestation in longitudinal placental studies.

A final, largely unaddressed variable when studying placental perfusion is how local injury alters blood delivery, both at term and across pregnancy. Pathological conditions have been known to alter placental perfusion [42,44], potentially affecting downstream physiological functions and fetal development. This is yet one more confounding factor to be considered in efforts to generalize the *ex vivo* model, which typically utilizes only healthy cotyledons, and should also be addressed for *in vivo* modeling. While healthy pregnancies often demonstrate a level of baseline pathology, increased injury may be necessary to initiate changes in maternal perfusion that lead to adverse outcomes observed clinically.

To this end, we developed a primate placental injury model to induce maternal vascular malperfusion lesions by injecting various dosages of Tisseel (Baxter International Inc., Deerfield, IL), a bioactive, cross-linked fibrin sealant used as hemostatic agent in vascular surgeries [45], directly into the IVS of the rhesus macaque placentae. Placental pathology is then assessed near full term using a novel quantitative approach [42], with basic injuries identified at high resolution in the cotyledons. With degrees of induced injuries quantified, the injury model can serve as an apparatus for evaluating relationships between tissue injuries and placental functions.

The aim of this study is therefore to incorporate these newly developed methods and address the unmet needs: We tracked placental perfusion domains across different gestational timepoints and measured their longitudinal perfusion changes in the rhesus macaque injury model with ferumoxytol DCE-MRI. The correspondence of MRI-derived perfusion domains and placental cotyledons was established by detailed comparisons with the cotyledons identified in the term placental tissue. The levels and degrees of pathology induced by the injury model compared to controls were further examined in a cotyledon-specific manner, and relationships between cotyledon perfusion and pathology were assessed within both control and Tisseel-treated placentae.

## Materials and Methods

The study design was approved by the Institutional Animal Care and Use Committee (IACUC) at our institution. In total, 13 rhesus macaques (RMs) were included in this study. Six RMs were injected with 0.5 mL (N=3; “Tisseel 1x”) or 1.5 mL (N=3; “Tisseel 3x”) of Tisseel into the anterior disc at gestational day (GD) ∼100 to induce various levels of placental injuries. As controls, 7 RMs were injected with 0.5 mL of saline into the amniotic fluid (N=3; “Saline AF”) at GD ∼55 or anterior placental disc (N=4; “Saline PL”) at GD ∼100. Both amniotic and placental injections were utilized to account for potential injury induced by direct injection into the placental tissue.

All subjects were imaged in right-lateral decubitus position on a 3T clinical MRI scanner (Discovery MR750 with a 32-channel torso coil/Signa Premier with a 30-channel air coil, GE Healthcare, Waukesha, WI) at up to 3 gestational timepoints, with the first scan on the day before injection, the second GD ∼115, and the third GD ∼145. The RMs were pre-medicated with ketamine (10 mg/kg) prior to the MRI scans, and general anesthesia with isoflurane gas was maintained during the scan [23], resulting in the sedation of both the mother and the fetus. Placental DCE-MRI data was acquired with ferumoxytol (Feraheme, 4 mg/kg, diluted 5:1 with saline) and a respiratory-gated, T1-weighted spoiled gradient echo sequence (DISCO, TR = 4.8 ms, TE = 1.8 ms, temporal resolution = 2.744 s, spatial resolution = 0.86x0.86x1.00 mm^3^, # of time frames=40). Ferumoxytol and a subsequent flush with 20 mL of saline were administered at 0.5 mL/s.

The RMs underwent cesarean section at GD ∼155 (full term = 165 days), at which point fetal and placental tissue were excised and weighed. Placental cotyledons were determined by visual and textural assessment, mapped onto whole-placental images, and dissected along septa. Center cuts of cotyledons were then obtained, paraformaldehyde-fixed, paraffin-embedded, stained with hematoxylin and eosin (H&E), scanned, and stitched together at 4x magnification [35]. Placental sub-tissue layers (chorionic plate, placental villi, intervillous space, and trophoblastic shell) and pathologies (fibrin deposition, villous agglutination, inflammatory agglutination, and stromal mineralization) were then digitally annotated using the GIMP (GNU Image Manipulation Program) software, with details described by Keding et al. [42]. Quantified cotyledon pathology was reported as the ratio of pathological area to total sub-tissue area and denoted as a percentage.

The MRI images were screened for phase wrap and motion artifacts to ensure image quality. Acquisitions with artifacts that were visually identifiable and present within the uterus were discarded from further processing. 3D segmentation of the placental boundary was performed with ITK-SNAP [46,47] based on images from the last point in the DCE time series by a graduate student with 3 years of experience (RYL) in segmenting rhesus macaque placental MRI images.

Flow and perfusion metrics were derived with a recently proposed workflow [39,40] and further optimized for noise reduction and flexibility of segmentation thresholds. Contrast arrival time *t_arr_* of each voxel was defined as the time corresponding to the maximum slope in the sigmoid fit of the temporal signal intensity (Figure 1). The volumetric additive inverse map of contrast arrival time for each placental disc (standard deviation = σ) was denoised by undergoing H-transform with thresholds of σ for local minima and 3σ for local maxima. Center and edge markers (representing early and late contrast arrival zones, respectively) were then generated by finding the local maxima and minima of the denoised and inversed arrival time map. 3D watershed segmentation (MATLAB R2025b, The Mathworks Inc., Natick, MA) was then performed on the inversed arrival time map with enhanced maxima/minima imposed on locations of center/edge markers to identify individual placental perfusion domains. The contrast-perfused volume-time function *V*(*t*) was defined as:

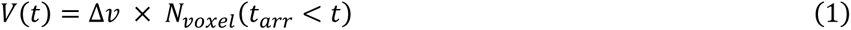

Where Δ*v* is the voxel size of image acquisition and *N_voxel_* is the number of voxel(s) satisfying the condition. For each perfusion domain, the blood flow was estimated using the maximum time derivative of the sigmoid fit of *V*(*t*), and the perfusion was calculated as flow divided by the domain volume.

**Figure 1.**
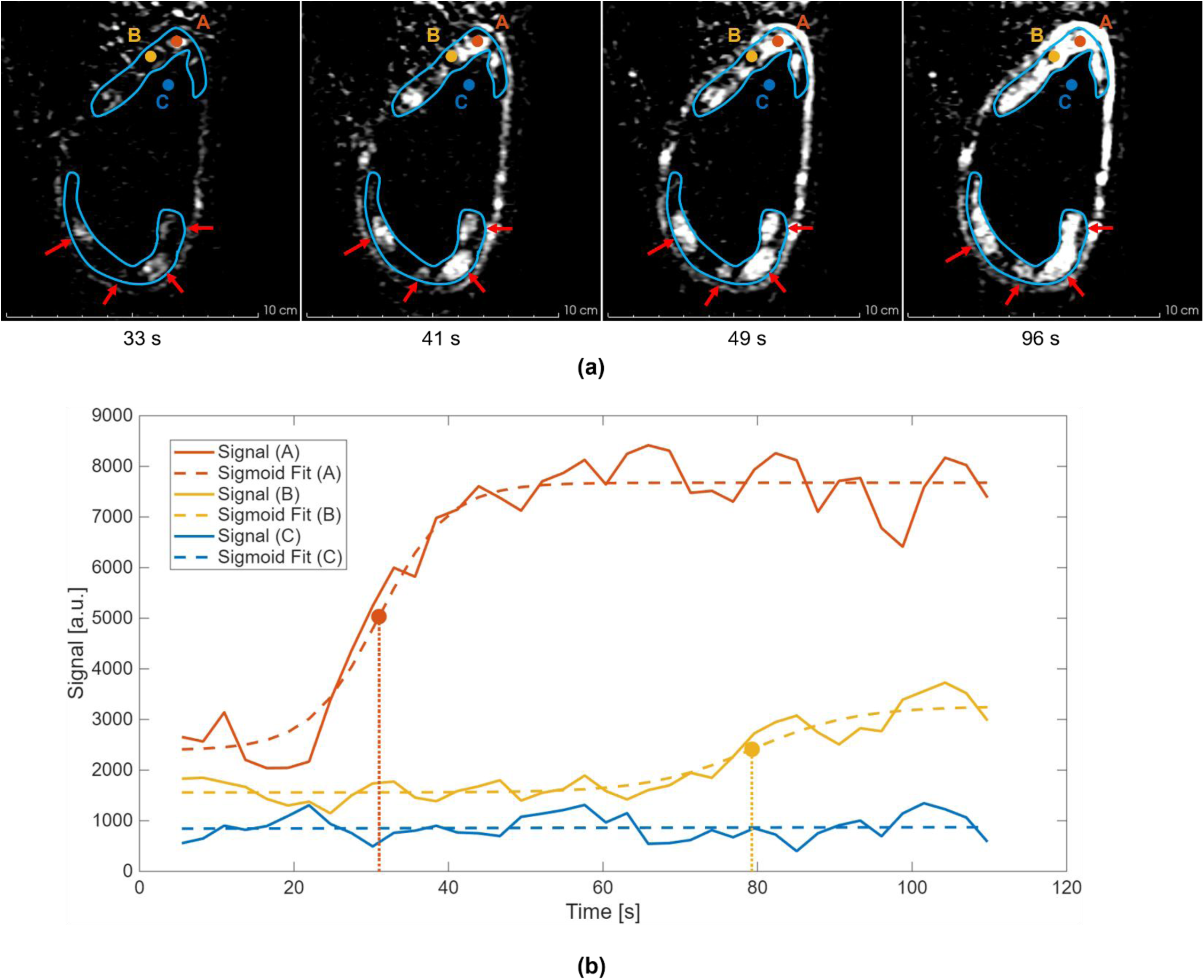
A representative example of contrast-enhanced time series images and voxel-based contrast arrival time determination. (a) Background-subtracted time series images of the placenta (blue lines) after ferumoxytol injection. The centers (e.g. point A) of perfusion domains (red arrows) enhance first, from where the contrast agent slowly advances into the peripheral areas (e.g. point B), demonstrating the perfusion behaviors in the placenta. The non-placental regions, including the amniotic fluid (e.g. point C) and the fetus, receive no ferumoxytol inflow. (b) The signal enhancement curves (solid lines) and their sigmoid fits (dash lines) of points A, B, and C, as denoted in (a). Within the placenta, the contrast arrival time of each voxel is defined by the corresponding time (dotted vertical lines) at the point of maximal slope (points A and B) on the signal sigmoid fit. The center of a perfusion domain (point A) has a smaller contrast arrival time and a larger signal intensity at the end of image acquisition compared to the peripheral area (point B).

The orientations of the placenta tissue and the virtual 3D placental volume were coaligned based on placental disc shape and the spatial patterns of septa and perfusion domain boundaries. In subjects where two placental discs were present, the disc correspondence of the tissue and virtual volume was established similarly. The cotyledons identified in the term placenta discs were then matched with one or more MRI-derived perfusion domain(s) identified at each gestational timepoint based on relative location, size, and boundary (RYL & LTK). Cotyledons not fully containing at least one perfusion domain were considered unmatched and excluded from further analyses.

All statistical analyses were conducted in MATLAB (R2025b). To evaluate the intergroup differences of quantified cotyledon pathology, each pathology marker first went through the nonparametric Kruskal-Wallis test [48] (α = 0.05) with the false discovery rate (FDR) controlled using the Benjamini-Hochberg procedure [49]. Markers showing significant general inter-group differences then underwent post hoc Dunn’s tests with Dunn–Šidák adjustment for pairwise group comparisons. To detect significant monotonous cotyledon perfusion changes over gestation, Theil-Sen (TS) and ordinary least square (OLS) slopes were calculated for the cotyledon perfusion versus gestation days and underwent two-tailed Wilcoxon signed-rank test (α = 0.05) based on treatment groups. The TS slopes were calculated in addition to the OLS slopes to account for potential outlier effects, and the Wilcoxon test was used instead of the t-test to generalize beyond the normal distribution assumption.

To evaluate monotonic correlations of all quantified pathology indicators versus both last-timepoint cotyledon perfusion and longitudinal cotyledon perfusion trends (TS and OLS slopes), Kendall’s tau and Spearman’s rho coefficients and their corresponding p-values were calculated (α = 0.05). Similarly, Kendall’s tau was calculated in addition to Spearman’s rho to account for potential outlier effects. In cases where no statistical difference was observed, power analyses for the statistical models used were performed with simulations (Number of permutations = 1e4, α = 0.05; test-specific parameters determined based on experiment results) to determine the least detectable effects with a statistical power (1 – β) = 80%.

## Results

Fetuses and dams from all Tisseel- and saline-treated pregnancies survived the total duration of experimentation. Of the 13 RMs, 10 received scans at all 3 gestational timepoints, while the other 3 RMs received scans at 2 timepoints. One scan of the latter showed motion artifacts and was excluded, resulting in 35 analyzed scans in total. Placentae and fetuses were successfully obtained from all C-section deliveries with weights recorded (Supplemental Table 1).

From the 13 placentae (23 placental total discs [10 bidiscoid, 3 monodiscoid]), 154 of 159 cotyledons were collected and received histopathological evaluation (Figure 2). The results of all pathological markers assessed within sub-tissues are summarized in Supplemental Table 2 and Supplemental Figure 1. Of the 20 distinct pathology and sub-tissue combinations, 17 were significantly elevated in the Tisseel-treated groups compared to saline-treated groups (Table 1). The Tisseel 1x group showed the highest percent inflammation and agglutination, whereas the Tisseel 3x group showed the highest percent mineralization. No significantly increased levels of pathology were observed in the Saline PL group compared to the Saline AF group.

**Figure 2.**
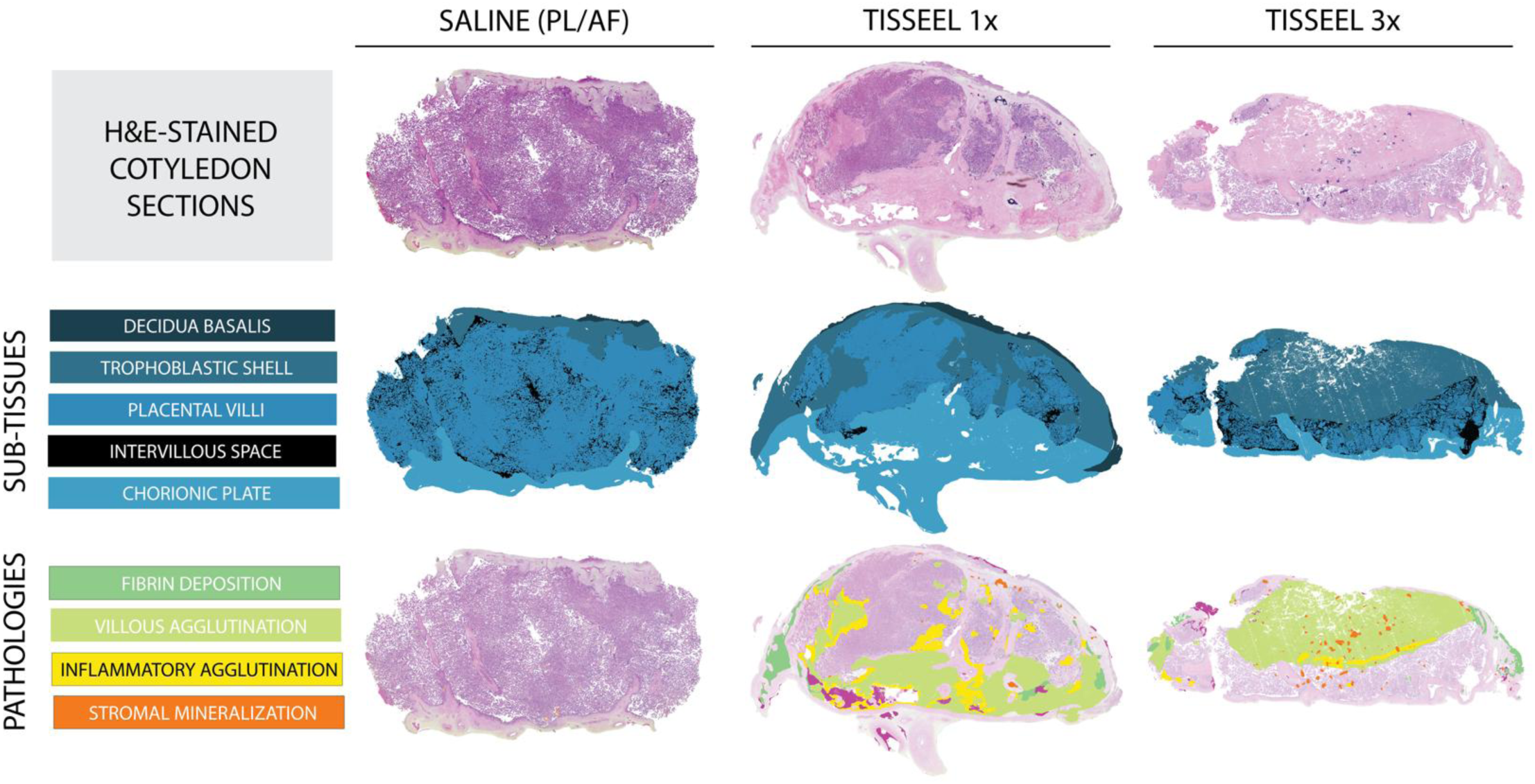
Representative annotated cotyledon slices from saline- and Tisseel-treated pregnancies. Tissue slices centrally cut through each cotyledon were stained with hematoxylin and eosin (H&E), then digitally annotated for sub-tissues and pathologies. Qualitative assessment revealed that cotyledons from Tisseel-treated pregnancies more often exhibited large, continuous pathological regions compared to cotyledons from saline-treated pregnancies. Saline AF: saline injected into amniotic fluid, Saline PL: saline injected into the placenta, Tisseel 1x: Tisseel 0.5 mL, Tisseel 3x: Tisseel 1.5 mL.

**Table 1.** Summary of pairwise intergroup pathology differences.

|  |  | P-Values | FDR Test | Significant Intergroup Difference |
| --- | --- | --- | --- | --- |
| **FBD** | **TOT** | 0.005* | Pass | T1 > (SP, SA) |
|  | **CHR** | 0.029* | Pass | None |
|  | **VIL** | 0.001* | Pass | SA > (T1, T3, SP) |
|  | **TBS** | 0.002* | Pass | (T1, T3) > SA |
| **AGL** | **TOT** | 0.001* | Pass | T1 > (T3, SP, SA) |
|  | **CHR** | < 0.001* | Pass | T1 > (T3, SP, SA) |
|  | **VIL** | 0.106 | Fail | -- |
|  | **TBS** | 0.002* | Pass | T1 > (T3, SP, SA) |
| **INF** | **TOT** | < 0.001* | Pass | T1 > (SP, SA); T3 > SP |
|  | **CHR** | < 0.001* | Pass | T1 > (SP, SA); T3 > SP |
|  | **VIL** | 0.162 | Fail | -- |
|  | **TBS** | 0.001* | Pass | T1 > (SP, SA) |
| **MIN** | **TOT** | < 0.001* | Pass | T3 > (T1, SP, SA) |
|  | **CHR** | < 0.001* | Pass | T3 > (T1, SP, SA) |
|  | **VIL** | < 0.001* | Pass | T3 > (T1, SP, SA) |
|  | **TBS** | < 0.001* | Pass | T3 > (T1, SP) |
| **PATH** | **TOT** | < 0.001* | Pass | (T1, T3) > (SP, SA) |
|  | **CHR** | < 0.001* | Pass | T1 > (SP, SA); T3 > SP |
|  | **VIL** | < 0.001* | Pass | T3 > (T1, SP, SA) |
|  | **TBS** | < 0.001* | Pass | (T1, T3) > (SP, SA) |

**Note:** Group X > (Group Y, Group Z): significantly higher pathology values in Group X than in Groups Y and Z. “None”: no significant pairwise intergroup difference.
**Abbreviations:** FDR: false discovery rate. SA: saline injected into amniotic fluid (“Saline AF”), SP: saline injected into the placenta (“Saline PL”), T1: Tisseel 0.5 mL (“Tisseel 1x”), T3: Tisseel 1.5 mL (“Tisseel 3x”). FBD: fibrin deposition, AGL: agglutination, INF: inflammation, MIN: mineralization, PATH: total pathology. TOT: whole slice, CHR: chorion, VIL: villus, TBS: trophoblast.

Each cotyledon identified in the term placental tissue was compared with the corresponding MRI-identified perfusion domain(s), demonstrating consistency across gestational timepoints (Figure 3). Out of the 159 cotyledons, 110 matched with exactly 1 perfusion domain at all timepoints (69.2%), while 13 cotyledons matched with 2 domains at all timepoints (8.2%), and 4 cotyledons matched with 3 domains at all timepoints (2.5%). In addition, 2 cotyledons (1.3%) did not match with any perfusion domain. The other 30 cotyledons (18.9%) corresponded to various numbers of domains at different timepoints (maximum variation = 2; max/min number of domains corresponding to one cotyledon = 4/1), among which 14 cotyledons matched with one perfusion domain at 2 out of 3 imaging timepoints (Figure 4).

**Figure 3.**
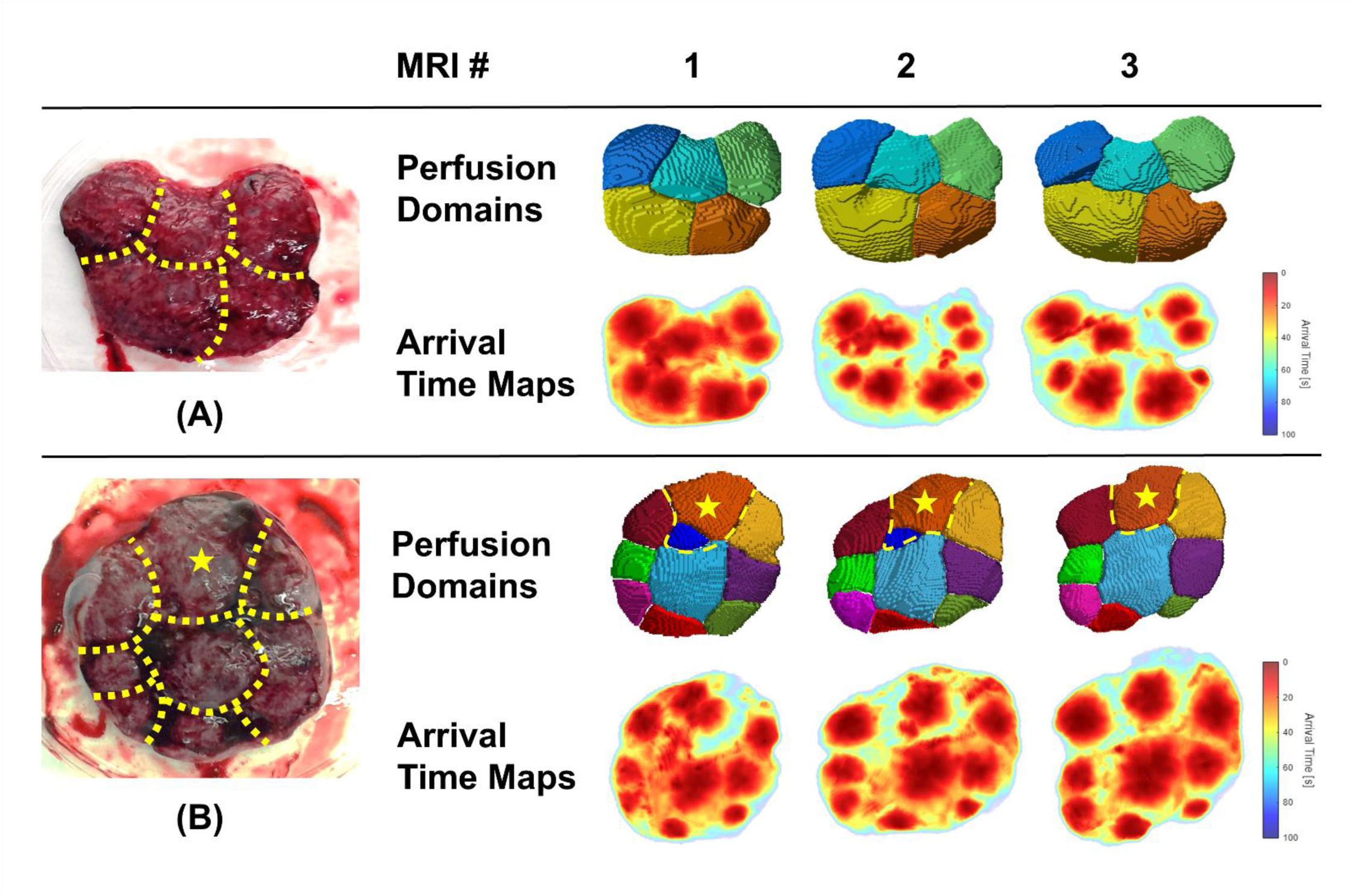
Two representative examples of term rhesus macaque placentae, perfusion domains, and arrival time maps of the same placental discs at all 3 gestational time points. Septa that separate individual cotyledons are marked with yellow dotted lines. The patterns of both arrival time maps and perfusion domains are consistent with the cotyledon pattern and over the observed period. All but one cotyledon in examples (A) and (B) match with exactly 1 perfusion domain at all timepoints; the star-marked cotyledon corresponds to 2/2/1 perfusion domains at timepoints 1/2/3, respectively.

**Figure 4.**
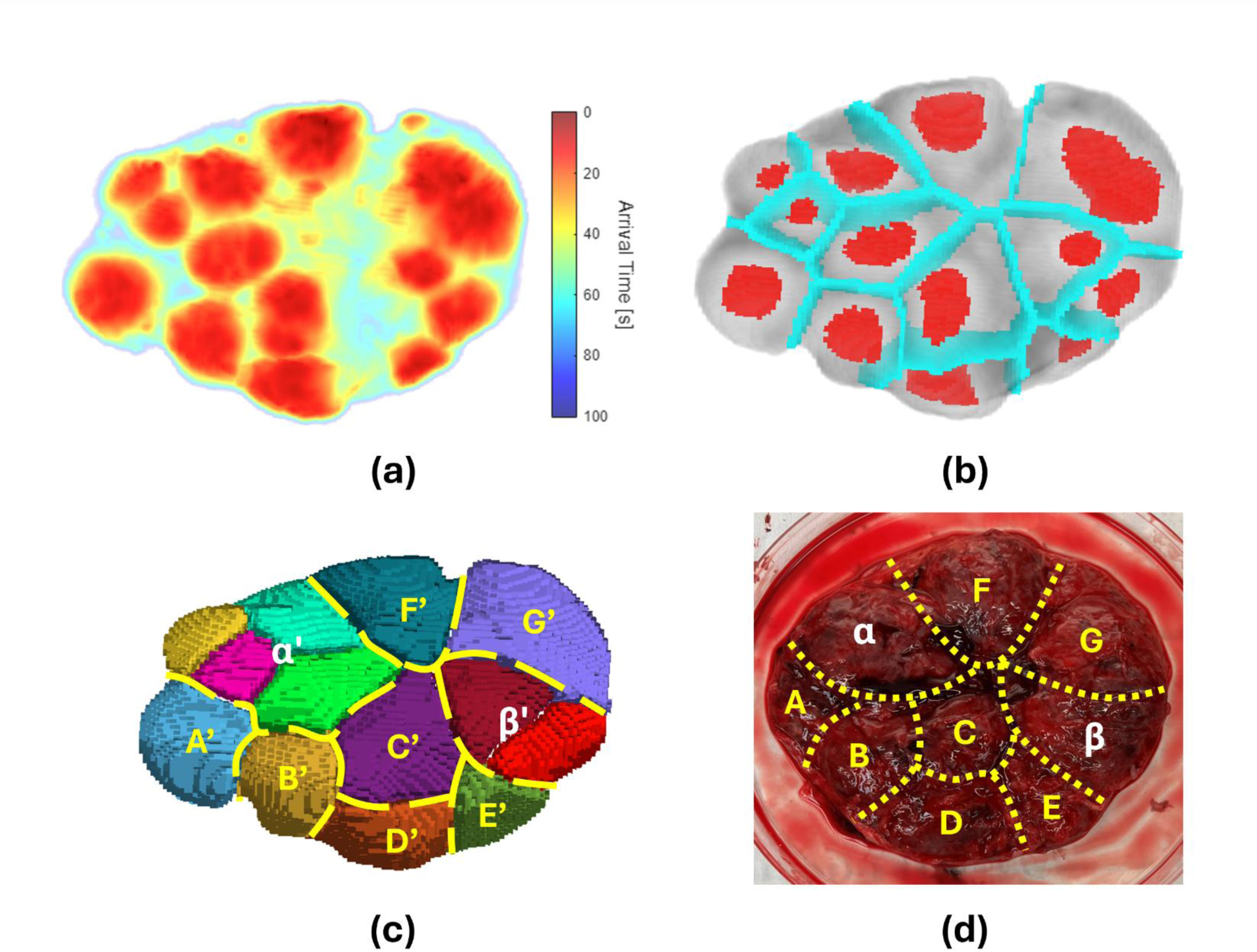
A representative example of contrast arrival time map, segmentation markers, perfusion domains, and cotyledons of the same rhesus macaque placental disc. (a) Contrast arrival time map. (b) Center (red) and edge (blue) guiding markers for watershed segmentation of perfusion domains. (c) Perfusion domains calculated by marker-guided watershed segmentation. (d) Term placental disc and observer identified septa (yellow dotted lines) that separate individual cotyledons. The septal pattern is consistent with the boundaries (yellow dashed lines) of perfusion domains in (c). Of the 9 cotyledons, 7 (A-G) have exactly 1-to-1 correspondence to the perfusion domains (A’-G’) in (c), while the remaining 2 cotyledons (α and β) correspond to 4 and 2 perfusion domains, respectively.

Across both saline- and Tisseel-treated groups, cotyledon volume and flow showed longitudinal increases (Figure 5). While the Saline PL group showed significantly decreased cotyledon perfusion across gestation, the Saline AF, Tisseel 1x, and Tisseel 3x groups showed no statistically significant relation between cotyledon perfusion and gestational days (Table 2). Simulation-based power analysis (30 cotyledons per group; standard deviation of between-subject variability = 0.005 mL/min/cc/GD; standard deviation of Gaussian measurement noise = 0.005 mL/min/cc/GD; probability of missing each scan = 10%) indicated ±0.0023 mL/min/cc/GD as the least detectable effect for both TS and OLS slopes using the study design and the statistical model (1 – β = 80%).

**Figure 5.**
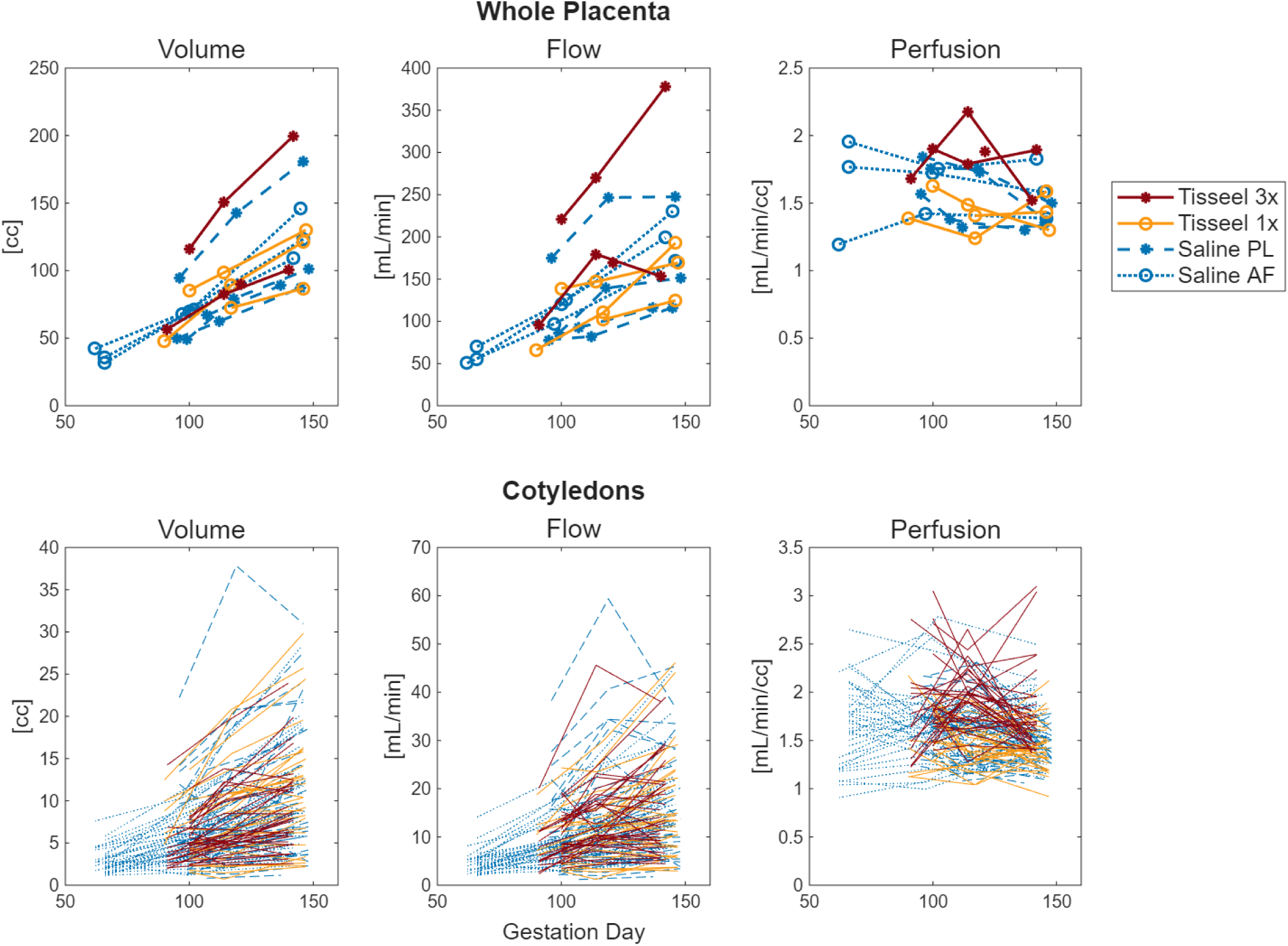
Longitudinal trends of volume, flow, and perfusion for whole placentae and cotyledons. For subjects with 2 placental discs, volume and flow were first calculated for each disc and then summed to calculate whole-placental perfusion. Saline AF: saline injected into amniotic fluid, Saline PL: saline injected into the placenta, Tisseel 1x: Tisseel 0.5 mL, Tisseel 3x: Tisseel 1.5 mL.

**Table 2.** Theil-Sen and ordinary least-squares slopes as measures of linear longitudinal perfusion changes and the corresponding two-tailed Wilcoxon signed-rank test results.

| [mL/min/cc/GD] | TS Slope |  |  | OLS Slope |  |  |
| --- | --- | --- | --- | --- | --- | --- |
|  | Mean | Median | P-Value | Mean | Median | P-Value |
| **Saline AF** | 0.0047 | 0.0015 | 0.259 | 0.0045 | 0.0016 | 0.282 |
| **Saline PL** | -0.0053 | -0.0038 | < 0.001* | -0.0051 | -0.0034 | < 0.001* |
| **Tisseel 1x** | -0.0010 | -0.0012 | 0.721 | -0.0009 | -0.0019 | 0.737 |
| **Tisseel 3x** | -0.0006 | -0.0036 | 0.523 | -0.0006 | -0.0033 | 0.799 |

**Abbreviations:** TS: Theil-Sen, OLS: ordinary least-squares. Saline AF: saline injected into amniotic fluid, Saline PL: saline injected into the placenta, Tisseel 1x: Tisseel 0.5 mL, Tisseel 3x: Tisseel 1.5 mL.

Finally, to determine whether increased injury uncovered significant pathology-perfusion relationships, correlations of cotyledon perfusion metrics versus pathological markers were evaluated in both saline- and Tisseel-treated groups. Of the total 159, 153 cotyledon pathological annotations were successfully paired with perfusion domains. Last-timepoint cotyledon perfusion showed significant positive correlations with agglutination and total pathology in villi (p < 0.05; Table 3). Longitudinal cotyledon perfusion trends (TS and OLS slopes) also demonstrated positive relationships with agglutination within the villous region, the trophoblastic shell, and across all sub-tissues (p < 0.05; Table 4). All pathology-perfusion correlation test results were consistent between Spearman’s/Kendall’s tests and TS/OLS slopes.

**Table 3.**
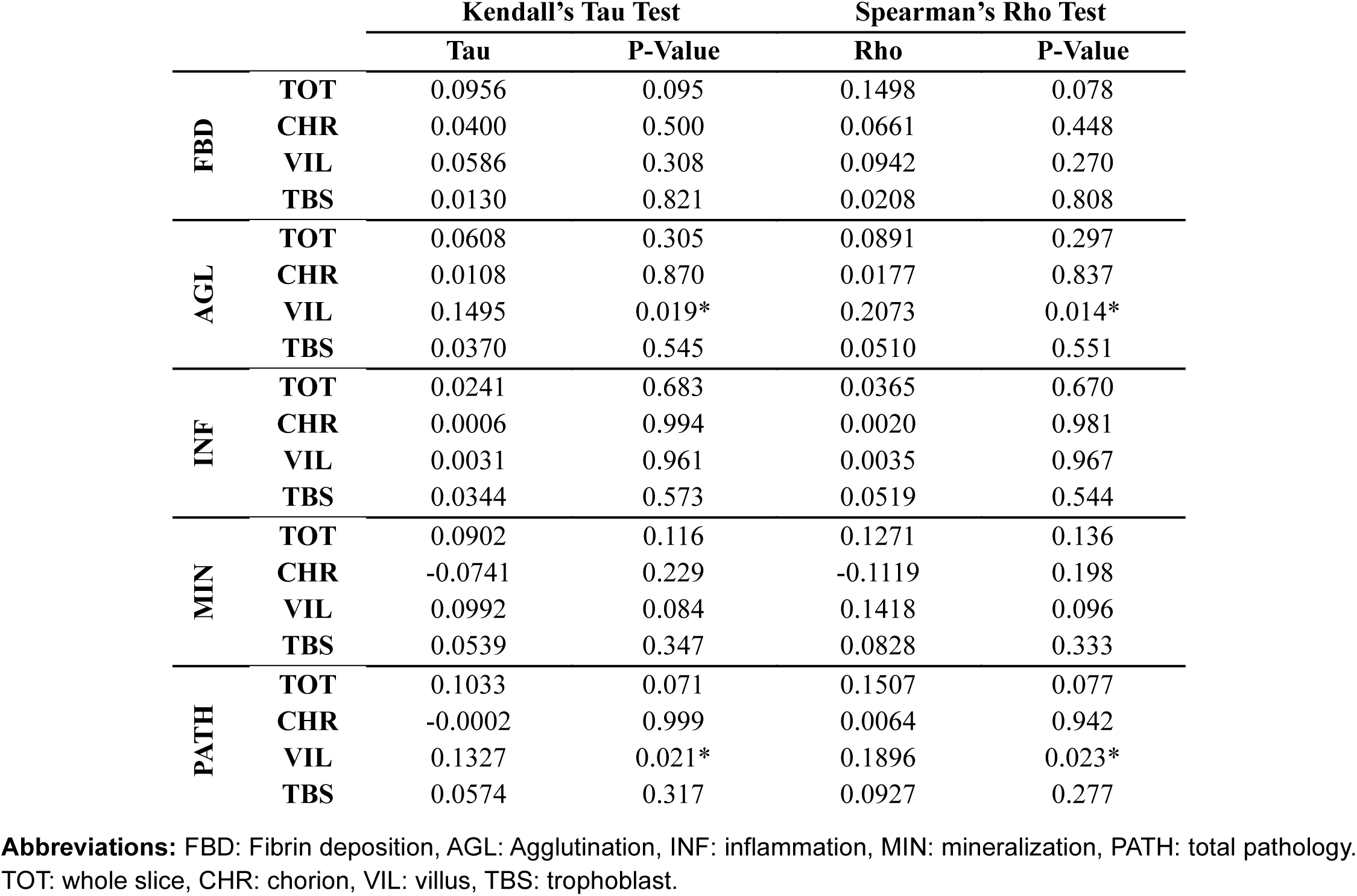
Correlation test results of last-timepoint cotyledon perfusion versus pathology metrics.

**Table 4.** Correlation test results of longitudinal perfusion changes versus pathology metrics.

|  |  | TS Slope |  |  |  | OLS Slope |  |  |  |
| --- | --- | --- | --- | --- | --- | --- | --- | --- | --- |
|  |  | Kendall's Tau Test |  | Spearman's Rho Test |  | Kendall's Tau Test |  | Spearman's Rho Test |  |
|  |  | Tau | P-Value | Rho | P-Value | Tau | P-Value | Rho | P-Value |
| **FBD** | **TOT** | 0.0709 | 0.248 | 0.1142 | 0.210 | 0.0820 | 0.181 | 0.1305 | 0.152 |
|  | **CHR** | -0.0068 | 0.916 | 0.0004 | 0.997 | -0.0021 | 0.976 | 0.0071 | 0.939 |
|  | **VIL** | 0.0450 | 0.465 | 0.0643 | 0.481 | 0.0490 | 0.426 | 0.0718 | 0.432 |
|  | **TBS** | -0.0278 | 0.652 | -0.0222 | 0.808 | -0.0115 | 0.853 | 0.0001 | 0.999 |
| **AGL** | **TOT** | 0.1735 | 0.006* | 0.2478 | 0.006* | 0.1698 | 0.007* | 0.2426 | 0.007* |
|  | **CHR** | 0.0956 | 0.172 | 0.1304 | 0.159 | 0.0867 | 0.215 | 0.1196 | 0.197 |
|  | **VIL** | 0.2350 | 0.001* | 0.3133 | 0.000* | 0.2315 | 0.001* | 0.3113 | 0.001* |
|  | **TBS** | 0.1527 | 0.019* | 0.2134 | 0.018* | 0.1590 | 0.015* | 0.2206 | 0.015* |
| **INF** | **TOT** | 0.0221 | 0.727 | 0.0388 | 0.671 | 0.0060 | 0.925 | 0.0174 | 0.849 |
|  | **CHR** | 0.0310 | 0.662 | 0.0451 | 0.628 | 0.0217 | 0.760 | 0.0277 | 0.766 |
|  | **VIL** | 0.0266 | 0.684 | 0.0340 | 0.710 | 0.0050 | 0.941 | 0.0036 | 0.969 |
|  | **TBS** | 0.0737 | 0.257 | 0.1031 | 0.258 | 0.0588 | 0.366 | 0.0843 | 0.356 |
| **MIN** | **TOT** | 0.1020 | 0.096 | 0.1602 | 0.078 | 0.0915 | 0.136 | 0.1381 | 0.129 |
|  | **CHR** | 0.0904 | 0.173 | 0.1306 | 0.159 | 0.0691 | 0.298 | 0.1045 | 0.260 |
|  | **VIL** | 0.0972 | 0.113 | 0.1425 | 0.117 | 0.0937 | 0.127 | 0.1376 | 0.131 |
|  | **TBS** | 0.0980 | 0.110 | 0.1561 | 0.086 | 0.0815 | 0.184 | 0.1249 | 0.170 |
| **PATH** | **TOT** | 0.1416 | 0.021* | 0.2178 | 0.016* | 0.1419 | 0.021* | 0.2110 | 0.020* |
|  | **CHR** | 0.0866 | 0.166 | 0.1415 | 0.126 | 0.0860 | 0.169 | 0.1358 | 0.143 |
|  | **VIL** | 0.1595 | 0.009* | 0.2266 | 0.012* | 0.1587 | 0.010* | 0.2218 | 0.014* |
|  | **TBS** | 0.0939 | 0.126 | 0.1523 | 0.094 | 0.0985 | 0.108 | 0.1552 | 0.088 |

**Abbreviations:** TS: Theil-Sen, OLS: ordinary least-squares. FBD: Fibrin deposition, AGL: Agglutination, INF: inflammation, MIN: mineralization, PATH: total pathology. TOT: whole slice, CHR: chorion, VIL: villus, TBS: trophoblast.

## Discussion

Currently, the connection between longitudinal maternal blood perfusion and significant placental injury is largely unknown. In this work, we successfully identified and tracked placental cotyledons at different gestational timepoints by meticulously matching perfusion domains derived by ferumoxytol DCE-MRI to term placental tissue. We further performed detailed digital annotation of cotyledons, demonstrating the ability of Tisseel injections at mid-gestation to induce higher levels of common placental pathologies near term compared to the saline-injected controls, supporting the efficacy of Tisseel injection to induce placental injury. Furthermore, by enhancing the sensitivity of the watershed segmentation workflow to early- and late-perfused areas with the application of segmentation markers and applying thorough comparisons across gestation timepoints, we were able to illustrate the feasibility of non-invasively tracking longitudinal perfusion changes in rhesus macaque cotyledons with various degrees of pathology. Importantly, we have continued to advance the resolution of pathological evaluation, from qualitative impression [40] or estimated pathological scoring [39] to quantitative, location-based pathological reporting.

The majority (69.2%) of the cotyledons identified in term placentae consistently matched with exactly one MRI-identified perfusion domain. It is noteworthy that another 10.7% of the cotyledons matched with 2 or 3 perfusion domains at all timepoints, and visual inspections of arrival time maps also confirmed the consistent multi-inflow nature of these cotyledons over time, further confirming the similar observation at a previous cross-sectional study [36]. This suggests that the contrast kinetics-based MRI approach on can potentially identify individual spiral artery inputs within cotyledons, providing a finer level of identifying placental function with *in vivo* imaging. Remarkably, variations of up to 2 in perfusion domain numbers at different timepoints were observed in 18.9% of the cotyledons, and similar phenomena were also previously observed in rhesus macaque placentae using x-ray angiography [50] and human placentae using blood oxygen level-dependent MRI [51]. This may reflect the temporary narrowing or occlusion of spiral arteries due to spontaneous vasoconstriction, of which the causes, extent, and frequency are still yet to be understood [52].

In our prior work focusing on healthy rhesus macaque pregnancy [42], we observed significant increases in compensatory perfusion with higher levels of stromal mineralization at term. Through increased placental injury induced by Tisseel injections, we further demonstrated both positive correlations of last-timepoint perfusion and longitudinal perfusion changes with villous agglutination. Inter-group comparisons also showed consistent results; for example, the Saline PL group demonstrated the lowest overall degree of pathology while being the only group with significantly decreased perfusion over time. Based on these data, it may be hypothesized that 1) maternal blood flow input exhibited compensatory increases in response to placental injury that outcompeted villous growth, and/or 2) villous growth was proportionally diminished in comparison to maternal blood flow in highly pathological cotyledons. The observed connection between pathology and perfusion in the primate placenta may have implications in the way we set out to treat or prevent placental injuries in humans.

There are limitations to this study. The number of subjects is small, due in part to long gestation periods of rhesus macaques, limited availability of research subjects, and high costs, thus limiting the statistical power of this work. Despite this, the cotyledon-specific resolution of this study greatly increased our ability to demonstrate precise injury with altered perfusion and allowed for strong statistical conclusions to be made. Simultaneously, the cohort size is appropriate to prove the feasibility of the approach and motivate future studies in animals and human subjects that benefit from the tracking of cotyledons and their individual function through pregnancy, enabled by non-invasive imaging. In addition, the non-parametric DCE-MRI analysis model used in this work is simplified compared to the complexity of anatomy and physiology in the placenta, yet it avoids the necessity of making excessive assumptions of tissue modeling and can thus be easily integrated into clinical applications where various levels of tissue pathology may be present.

Further investigations on the tolerance of the maximal safe ferumoxytol infusion rate and the minimal necessary dosage based on required MR contrast and signal-to-noise ratios are needed for the translation of the protocol in this study into human placental research and clinical practice. The anatomical and physiological differences between human and rhesus placentae, especially the depth of septa reaching into the intervillous space [53], should also be taken into account as it can affect the perfusion behaviors of individual cotyledons. Cotyledon tissue or MR-compatible perfusion phantoms that closely resemble human cotyledon hemodynamics [54] may therefore be used in *ex vivo* experiments to determine appropriate ferumoxytol dosages for detecting separate spiral artery entries. With these constraints established, placental functions in and interactions of individual cotyledons may be more thoroughly studied.

## Conclusion

This study demonstrates the feasibility of noninvasively identifying placental cotyledons and longitudinally tracking cotyledon perfusion using ferumoxytol DCE-MRI, as well as confirms the efficacy of the rhesus macaque thrombotic injury model. The observed positive correlations of cotyledon perfusion and quantified pathology suggest the presence of intrinsic placental regulatory mechanisms and functional plasticity. With further optimization of imaging protocols and consideration of interspecies differences, this research framework could help validate *ex vivo* perfusion models and improve assessment of placental functions across gestation.

## Acknowledgements

The study was supported by the National Institutes of Health Grants R01-HD103443 and R01-AI132519. JV is supported by K01-AI182448 and LTK is supported by T32-HD101384. The Wisconsin National Primate Research Center is supported by P51-OD011106 and C06-RR020141. The content of this manuscript is solely the responsibility of the authors and does not represent the official views of the NIH. We gratefully acknowledge GE Healthcare for research support of University of Wisconsin-Madison, and AMAG Pharmaceuticals for providing ferumoxytol used for this study. We also thank the Wisconsin National Primate Research Center Veterinary, Scientific Protocol Implementation, and Animal Services staff for providing animal care, and assisting in procedures including breeding, pregnancy monitoring, and sample collection.

---

**Supplemental Table 1.** Fetal and placental weights.

| Treatment | Subject | Fetal Weight [g] | Placental Weight (Disc 1) [g] | Placental Weight (Disc 2) [g] | Total Placental Weight [g] | Fetal/Placental weight ratio |
| --- | --- | --- | --- | --- | --- | --- |
| Saline AF | r02034 | 504.0 | 72.7 | 62.9 | 135.7 | 3.72 |
|  | r12036 | 575.0 | 74.3 | 42.7 | 117.0 | 4.92 |
|  | rh2804 | 530.0 | 47.7 | 47.6 | 95.3 | 5.56 |
| Saline PL | rh3014 | 490.0 | 88.9 | 31.0 | 119.8 | 4.09 |
|  | rh3015 | 420.0 | 45.3 | 49.2 | 94.5 | 4.45 |
|  | rh2936 | 471.0 | 53.1 | 41.9 | 95.0 | 4.96 |
|  | rh2522 | 550.0 | 139.1 | -- | 139.1 | 3.95 |
| Tisseel 1x | rh3014 | 395.0 | 128.8 | -- | 128.8 | 3.07 |
|  | rh3051 | 450.0 | 97.5 | -- | 97.5 | 4.62 |
|  | r05055 | 478.0 | 67.2 | 59.0 | 126.2 | 3.79 |
| Tisseel 3x | rh3015 | 400.4 | 57.3 | 51.8 | 109.1 | 3.67 |
|  | r13088 | 470.0 | 62.7 | 63.1 | 125.8 | 3.74 |
|  | r05012 | 622.7 | 121.3 | 90.5 | 211.7 | 2.94 |

**Supplemental Table 2.** Pathology marker percentages in different tissues.

| [%] |  | Saline AF |  |  | Saline PL |  |  | Tisseel 1x |  |  | Tisseel 3x |  |  |
| --- | --- | --- | --- | --- | --- | --- | --- | --- | --- | --- | --- | --- | --- |
|  |  | Mean | Median | SD | Mean | Median | SD | Mean | Median | SD | Mean | Median | SD |
| **FBD** | **TOT** | 1.36 | 0.76 | 2.79 | 1.57 | 0.54 | 3.54 | 2.61 | 2.02 | 2.34 | 2.57 | 1.11 | 4.63 |
|  | **CHR** | 1.35 | 0.29 | 2.15 | 0.66 | 0.28 | 0.88 | 2.17 | 1.14 | 2.91 | 4.02 | 0.80 | 6.60 |
|  | **VIL** | 0.99 | 0.26 | 4.04 | 1.02 | 0.12 | 5.25 | 0.52 | 0.08 | 1.97 | 0.18 | 0.09 | 0.29 |
|  | **TBS** | 2.69 | 1.89 | 2.65 | 4.24 | 2.46 | 5.25 | 5.97 | 4.82 | 4.94 | 6.57 | 4.61 | 6.96 |
| **AGL** | **TOT** | 0.67 | 0.17 | 1.24 | 0.78 | 0.17 | 1.60 | 7.30 | 2.21 | 11.63 | 3.07 | 0.27 | 8.67 |
|  | **CHR** | 1.23 | 0.00 | 3.12 | 0.77 | 0.00 | 2.18 | 10.59 | 2.88 | 16.34 | 3.79 | 0.00 | 10.32 |
|  | **VIL** | 0.06 | 0.00 | 0.18 | 0.11 | 0.00 | 0.36 | 0.26 | 0.00 | 0.70 | 0.21 | 0.00 | 0.93 |
|  | **TBS** | 2.04 | 0.00 | 3.45 | 2.21 | 0.00 | 3.33 | 11.47 | 5.91 | 15.15 | 5.08 | 0.01 | 12.98 |
| **INF** | **TOT** | 0.37 | 0.13 | 0.58 | 0.33 | 0.07 | 0.58 | 2.71 | 1.52 | 3.44 | 1.19 | 0.65 | 1.49 |
|  | **CHR** | 0.33 | 0.00 | 0.93 | 0.43 | 0.00 | 1.70 | 6.15 | 1.38 | 7.77 | 3.06 | 0.00 | 5.50 |
|  | **VIL** | 0.07 | 0.02 | 0.13 | 0.16 | 0.00 | 0.58 | 0.31 | 0.00 | 0.63 | 0.24 | 0.02 | 0.71 |
|  | **TBS** | 1.60 | 0.00 | 2.31 | 0.75 | 0.00 | 1.42 | 3.79 | 2.63 | 4.18 | 2.37 | 0.69 | 3.89 |
| **MIN** | **TOT** | 0.67 | 0.50 | 0.58 | 0.66 | 0.43 | 0.63 | 0.54 | 0.49 | 0.43 | 1.47 | 1.25 | 0.84 |
|  | **CHR** | 0.29 | 0.00 | 0.59 | 0.34 | 0.02 | 0.99 | 0.15 | 0.05 | 0.29 | 0.68 | 0.40 | 0.82 |
|  | **VIL** | 0.49 | 0.25 | 0.55 | 0.41 | 0.22 | 0.61 | 0.47 | 0.28 | 0.53 | 1.31 | 1.00 | 0.97 |
|  | **TBS** | 2.00 | 1.69 | 1.82 | 2.10 | 1.02 | 2.31 | 1.06 | 0.72 | 0.91 | 2.59 | 2.53 | 1.42 |
| **PATH** | **TOT** | 3.07 | 1.94 | 3.56 | 3.35 | 1.96 | 5.06 | 13.15 | 6.10 | 15.15 | 8.30 | 4.48 | 12.61 |
|  | **CHR** | 3.21 | 1.13 | 4.79 | 2.20 | 0.69 | 3.31 | 19.06 | 14.36 | 19.83 | 11.55 | 3.50 | 17.37 |
|  | **VIL** | 1.62 | 0.69 | 4.24 | 1.71 | 0.66 | 5.97 | 1.55 | 0.62 | 2.68 | 1.95 | 1.43 | 1.66 |
|  | **TBS** | 8.34 | 6.85 | 5.93 | 9.29 | 8.98 | 7.44 | 22.29 | 12.79 | 19.57 | 16.62 | 13.18 | 16.20 |

**Notes:** Cotyledons without certain tissue layer(s) being present in the slice were excluded from the corresponding tissue-specific analysis. The smaller medians compared to the means and the high standard deviations indicated highly positive skewness for the majority of pathological markers.
**Abbreviations:** SD: sample standard deviation. FBD: Fibrin deposition, AGL: Agglutination, INF: inflammation, MIN: mineralization, PATH: total pathology. TOT: whole slice, CHR: chorion, VIL: villus, TBS: trophoblast.

**Supplemental Figure 1.**
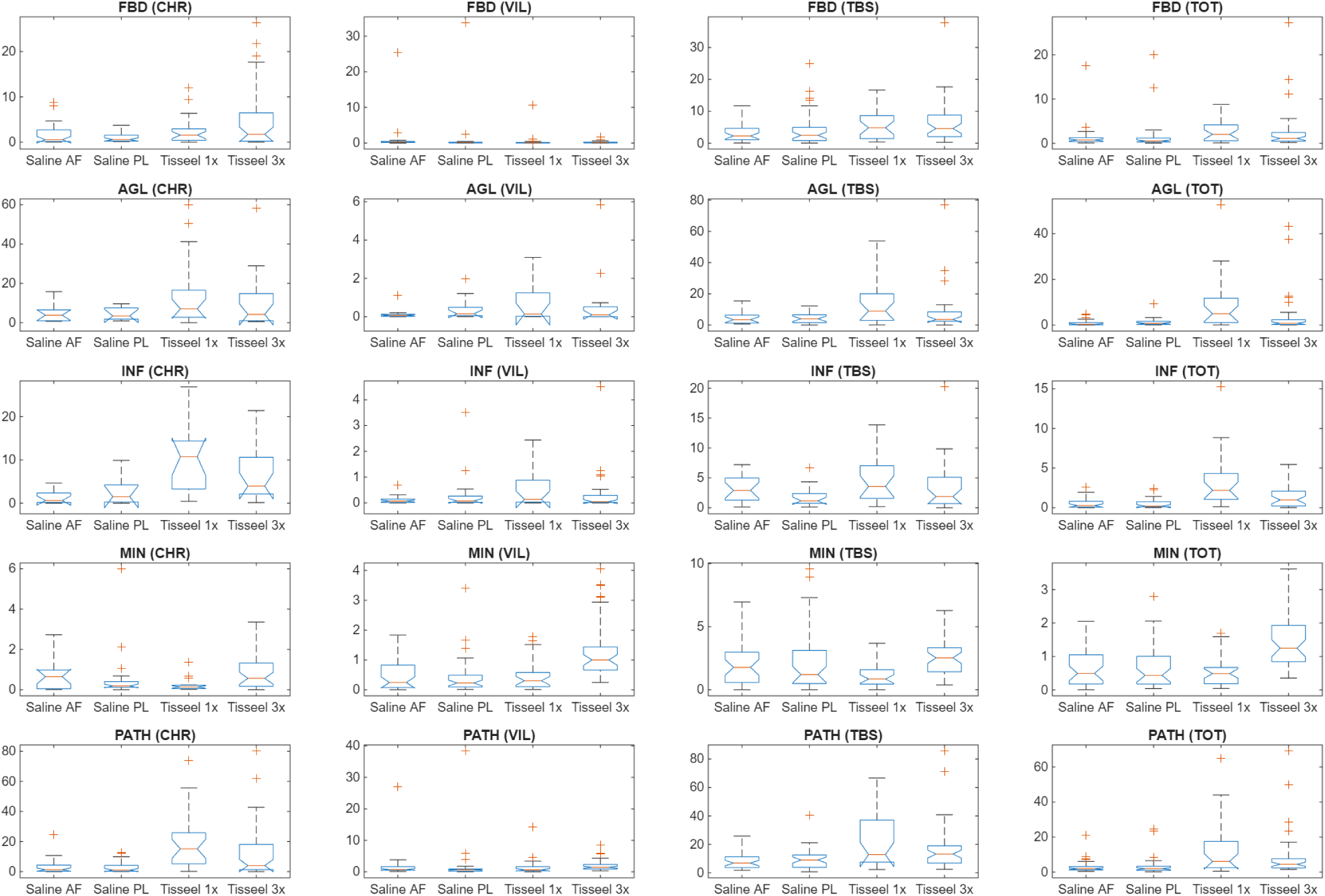
Cotyledon pathology percentages of all different pathology markers summarized using notched box plots. The vertical axes represent the percentages [%]. FBD: Fibrin deposition, AGL: Agglutination, INF: inflammation, MIN: mineralization, PATH: total pathology. CHR: chorion, VIL: villus, TBS: trophoblast, TOT: whole slice.

